# Why is the average collateral effect of synonymous mutations so similar across alternative reading frames?

**DOI:** 10.1101/2022.03.22.485379

**Authors:** Stefan Wichmann, Zachary Ardern

## Abstract

The standard genetic code has been shown to have multiple interesting properties which impact on molecular biology and the evolutionary process. One facet of molecular biology where code structure is particularly important is the origin and evolution of overlapping genes. We have previously reported that the structure of the standard genetic code ensures that synonymous mutations in a protein coding gene will lead to a remarkably similar average “collateral” mutation effect size in at least four out of the five alternative reading frames. Here we show that only 0.26% of alternative codes with the block structure of the standard genetic code perform at least as well as the standard code in this property. Considering this finding within a code optimality framework suggests that this consistent effect size across the different frames may be adaptive. Here we give context for this finding and present a simple model where a trade-off between evolvability and robustness leads to an average mutation effect size which maximises population fitness. This supports the intuition that similar mutation effects across the different alternative reading frames may be an adaptive property of the standard genetic code which facilitates evolvability through the use of alternative reading frames.

## Introduction

Protein coding in alternative reading frames of known genes was a topic of interest early in the development of modern molecular genetics, and has recently been receiving more attention. Overlapping genes (OLGs) were first found in bacteriophages (Barrell, Air, and Hutchison 1976), and among biologists they are often assumed to be restricted to viruses. There are indeed many prominent examples in viruses beyond bacteriophages including in the pandemic viruses HIV and SARS-CoV-2 (Firth and Brierley 2012; Cassan et al. 2016; Affram et al. 2019; Nelson et al. 2020; Firth 2020), but they have also recently been discovered in diverse cellular organisms, including bacteria (Kreitmeier et al. 2022; Zehentner et al. 2020b; Meydan, Vázquez-Laslop, and Mankin 2018), archaea (Gelsinger et al. 2020), and mammals, including humans (Loughran et al. 2020; Khan et al. 2020; Mudge et al. 2021; Cao et al. 2021; Wright et al. 2022). Research on OLGs within five years of their initial discovery investigated diverse topics including triple overlaps (Szekely 1978), information theory (Siegel and Fitch 1980; Smith and Waterman 1981; Yockey 1981), discussion of their evolution (Miyata and Yasunaga 1978; Yockey 1979), and the proposal that they may be widespread throughout life (Kolata 1977). The diverse evidence for overlapping genes accumulated in the decades since, along with their biological roles and potential biosynthetic applications, have recently been reviewed (Wright, Molloy, and Jaschke 2022).

Is there a functional reason for the maintenance of overlapping genes in genomes, or the use of alternative frame sequences in protein coding more generally? While overlapping genes have diverse individual functions, it is often thought that genome compression is the main function of overlapping genes as a class of genes. A study directly addressing this question has however suggested that the hypothesis of evolutionary pressure for smaller genomes doesn’t suffice as an explanation for OLG distribution, and in fact their role in evolution of functional novelty may be more important (Brandes and Linial 2016). Gene origin from non-coding sequences has received extensive attention recently, with a revolution in our understanding of protein evolution to incorporate many instances of genuinely ‘de novo’ origin (Vakirlis, Carvunis, and McLysaght 2020). Alternative frame sequences have previously been proposed as a source of evolutionarily novel genes, originating through a process called “overprinting” (Keese and Gibbs 1992). They were even mentioned in the prominent book by Susumu Ohno that developed the standard hypothesis that most genes arise through duplication and divergence from ancestral genes (Ohno 1970). This overprinting hypothesis has gained recent attention regarding the origins of genetic novelty (Carter 2021). A number of other papers on OLGs in bacteria have also been published (Hücker, Vanderhaeghen, Abellan-Schneyder, Scherer, et al. 2018; Hücker, Vanderhaeghen, Abellan-Schneyder, Wecko, et al. 2018; Vanderhaeghen et al. 2018; Zehentner et al. 2020a; Kreitmeier et al. 2022), although the focus has generally been on establishing their expression as proteins, and elucidating associated phenotypes, rather than their evolution or potential contribution to gene origins more broadly. Aside from overprinting, two related and previously unknown mechanisms for protein novelty from alternative reading frames - gene remodelling and pairs of compensatory frameshift mutations - have recently been elucidated (Watson, Lopez, and Bapteste 2022; Biba, Klink, and Bazykin 2022), as discussed later in this study.

An overlapping gene pair or alternative frame sequence locus consists of a reference frame sequence and alternative frame sequence (Figure 1). Elsewhere these have been referred to as the (pre-existing) “mother gene” and (younger, overprinted) “daughter gene” by analogy to mother and daughter cells in reproduction (Kreitmeier et al. 2022). Our target however is broader in also considering alternative frame sequences which are not functional genes, so we refer simply to reference and alternative reading frames. Some properties of the sequences in alternative reading frames are dictated by the structure of the standard genetic code. For a novel protein sequence encoded in an alternative frame of an existing protein-coding gene, the position in relation to the pre-existing ‘reference’ frame (i.e. relative reading frame) will determine some of its properties, given a particular genetic code mapping between codons and amino acids. For instance, the two alternative reading frames on the same strand as a given “reference frame” sequence tend to encode amino acids which preserve the hydrophobicity profile of the reference frame’s protein sequence (Bartonek, Braun, and Zagrovic 2020). This finding has been argued to not be surprising, as an artefact of possible selection on the code for robustness to point mutations (Xu and Zhang 2021a), so we have not listed it here as an ‘optimality’ of the code, but it may deserve further attention, as we have previously shown that both purported optimalities and claims for a lack thereof are sensitive to code sets, threshold choices, and the combination of properties used (Wichmann and Ardern 2019). The underlying assumption that frameshifts have no biological use is also called into question by some of the studies discussed here and our results. Aside from the tendency for preservation of hydrophobicity in same strand alternative reading frames, the structure of the standard genetic code also determines the hydrophobicity of the reading frame directly antisense (−1) to a reference sequence, with a strong tendency for amino acids of opposite hydrophobicities to be encoded by the codons in antisense to each other - and this property may assist in templating structural elements in antisense open reading frames (Konecny et al. 1993; Zull and Smith 1990). Not all properties of overlapping genes are simply a result of code structure, however; for instance the higher intrinsic disorder of overlapping versus non-overlapping proteins is not due to code structure (Willis and Masel 2018).

**Figure 1:**
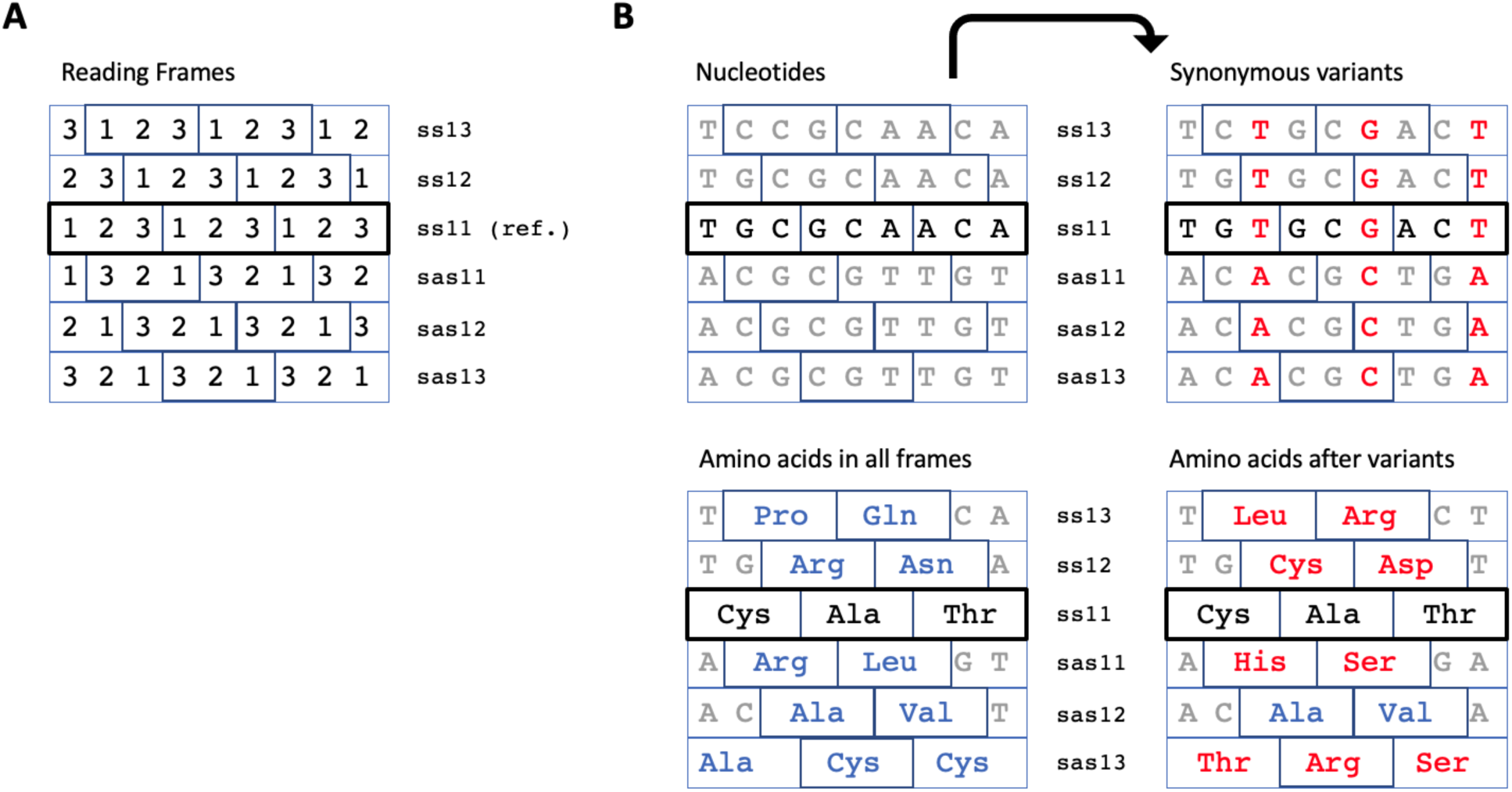
**A)** The five alternative reading frames, named according to (Wei and Zhang 2015; Nelson, Ardern, and Wei 2020) (ss=sense-sense; sas=sense-antisense; the two numbers represent the codon position in the reference and alternative frame respectively). **B)** Example of ‘collateral’ effects in alternative frames following synonymous mutations in the reference frame; variants at the nucleotide level and changes in amino acids encoded by alternative reading frames are shown in red - original amino acids are shown in blue.

The arrangement of the standard genetic code could very plausibly be different, resulting in different properties in alternative reading frames. Indeed, there are variant genetic codes across diverse taxonomic groups, although these minor variants extant today appear to all be derivative of the standard code (Osawa 1995; S. J. Freeland et al. 2000). A triplet code encoding at most the 20 canonical amino acids can be arranged in more than 10^80^ different ways. The actual arrangement has been shown to be somewhat optimal in comparison to alternative possible codes, across various biologically useful features, including robustness to point mutations (S. J. Freeland and Hurst 1998), termination after a frameshift, and the incorporation of additional non-coding information within protein-coding sequences (Itzkovitz and Alon 2007). We summarised some of this literature and analysed a few properties in a previous study (Wichmann and Ardern 2019). The choice of the 20 amino acids is also near optimal in its coverage of physico-chemical space (Ilardo et al. 2015, 2019; Mayer-Bacon and Freeland 2021). Intriguingly, the structure of the code is not just beneficial for biological functions such as minimising the effect of mistranslation, but also facilitates evolution, insofar as minimising mutation effect size promotes adaptation, as seen e.g. in Fisher’s Geometric Model of adaptation (Stephen J. Freeland 2002; Zhu and Freeland 2006). More strikingly, the standard code facilitates the exploration of sequence space better than if it merely reduced mutation effect size, as the code helps ensure that mutations are both depleted for deleterious variants and enriched for adaptive variants (Firnberg and Ostermeier 2013), this optimises the exploration of functional variants at intermediate time scales, supported by a study using a larger experimental fitness landscape (Tripathi and Deem 2018). While investigating the potential multi-dimensional optimality of the code inferred from the studies cited here, we previously discovered that synonymous mutations in a reference reading frame have remarkably similar average mutation effects (Wichmann and Ardern 2019). In this study we investigate potential ramifications of this finding further, by means of a simple evolutionary model.

Evolution by natural selection can be visualised as a process of climbing peaks in a fitness landscape. The topology of real fitness landscapes is still being investigated (Richter and Engelbrecht 2014; de Visser and Krug 2014; Payne and Wagner 2019; Chen, Fowler, and Tokuriki 2022), but it has been shown they are often ‘rugged’, meaning that there are local fitness peaks in addition to the global peak. Such ruggedness can limit adaptive evolution, depending on the height of the local peaks and their proximity to a global peak. Considerations of fitness landscapes in the context of individual genes generally concern situations where a functional sequence (e.g. a protein-coding gene) is undergoing adaptation. Here we are interested in processes of sequence exploration more generally, particularly the origin of new functional sequences. This requires not just minor adaptation, but also a wider exploration of sequence space - as such, we consider two different types of evolutionary events. The first are small effect mutations, which are more likely to shift a sequence towards a fitness peak, in line with standard Darwinian processes as seen for instance in Fisher’s Geometric Model (Tenaillon 2014; Fisher 1930). The second are large effect “evolvative” mutations, which assist with moving between fitness peaks (local maxima), preventing populations getting stuck in small local maxima, and more generally can sample disparate regions of sequence space as compared to local search.

## Methods

The methods for the calculation of alternative frame average mutation effect size in a given genetic code are reported in our previous study (Wichmann and Ardern 2019). Following a standard approach in the genetic code optimality literature (Buhrman et al. 2011), “mutation effect size” is measured as the square of the difference in polar requirement between two amino acids. The value of interest here is the average effect size in alternative reading frames of mutations which are synonymous in a reference reading frame. i.e. the average effect size in the amino acids encoded in alternative frame codons following synonymous mutations in reference reading frame codons. The “minus 1” frame (sas13 in Figure 1) was not included among the alternative frames, as it overlaps a single codon in the reference frame while other alternative frames overlap dicodons, and no method to directly compare them was found. The average mutation effect size in the four other alternative reading frames was calculated for all codons in a large set of 10^7^ possible genetic code tables sharing the block structure of the standard genetic code, including the standard genetic code itself.

The model in this study investigates whether an average mutation effect size (step size in our model) can be chosen so as to maximise the number of sequences ending up in a large fitness peak. We represent sequence space as a 2D surface with periodic boundary conditions (i.e. the space in the model loops back on itself so there is no ‘edge’ to the map). Particles represent sequences, which shift in sequence space over the course of generations, due to the effects of mutations (Figure 2A). Sequences are altered (particles move around the map) in one of two ways - in steps which are either ‘conservative’ or ‘evolvative’ (Figure 2B; Table 1). The ‘conservative’ mutations have a small step size *s*_*c*_ and have an inbuilt higher chance of moving the sequence to higher fitness. That is, if the sequence is within a fitness peak region, these mutations are biased towards moving towards the centre of the peak, modelling the influence of the filter of natural selection (positive selection) on such mutations. The direction of the larger ‘evolvative’ mutations on the other hand is not influenced by fitness, and as such they have a randomly chosen direction, i.e. all angles are equiprobable regardless of the particle’s location in the map. These represent mutations with a large effect which evade purging by natural selection.

**Table 1:** Summarised properties of the two types of evolutionary “step” within the model.

| Mutation type | Step size | Biased towards higher fitness? | Probability of this mutation |
| --- | --- | --- | --- |
| conservative | $s_c$ | Yes, if within a peak | $p_c$ |
| evolvative | $s_e$ | No | $1-p_c$ |

**Figure 2:**
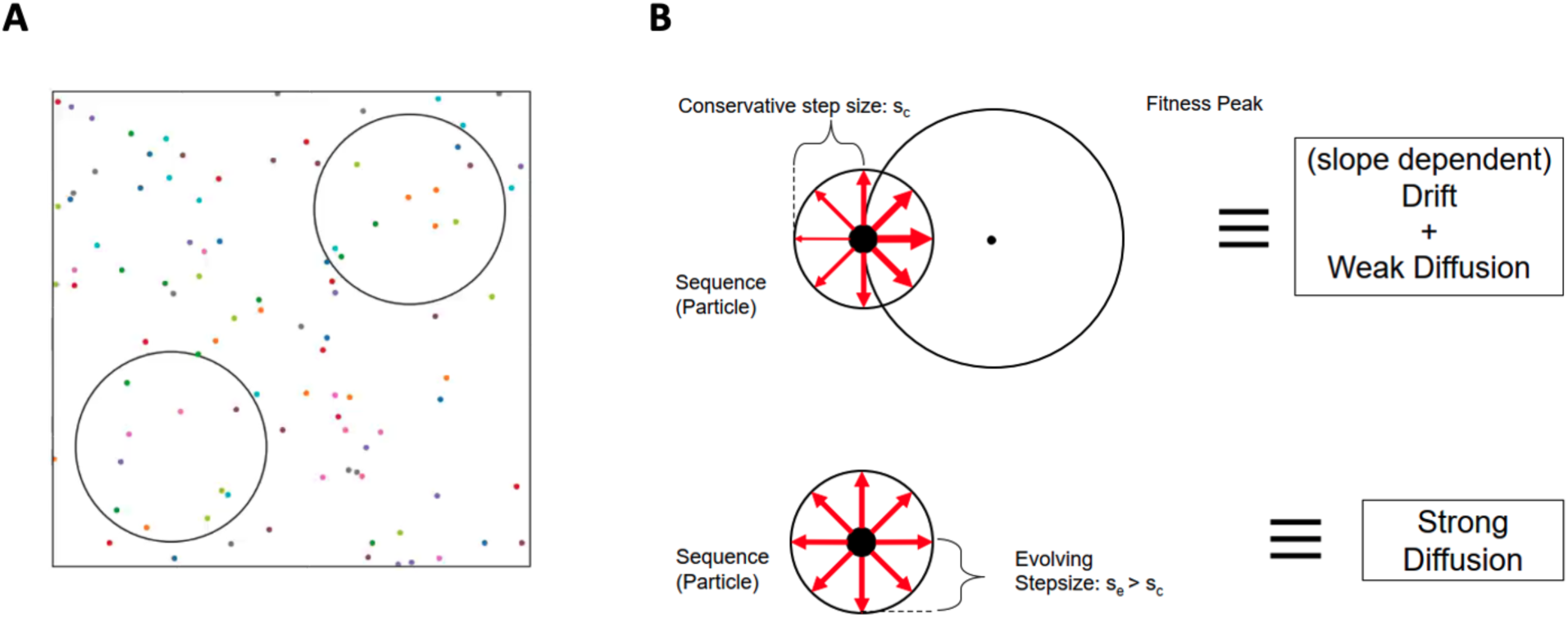
**A)** In this 2D representation of the model, sequence space is represented with circles corresponding to fitness peaks, and an initial distribution of sequences represented as points. Each fitness peak is a symmetric cone with radius 0.2 in a normalised sequence space of size 1×1. The top right peak’s fitness value (height, H) is 160 and for the lower left peak H=40. **B)** Illustration of conserving and evolving mutations in the model - the direction of conserving mutations is affected by whether the sequence affected is situated within a peak, while evolving mutations are not.

The direction of movement of the steps (of size *s*_*c*_ or *s*_*e*_) is random, but in the case of conservative mutations the probability of moving in each possible direction depends on the relative fitness for this direction as compared to others. In detail, the possible directions for mutational steps are discretized into N equiangular directions, with N=100 used throughout this study. The fitness *f*_*i*_ value after each possible step move is calculated, and from this the minimum fitness after a movement, *f*_*min*_, is determined. The probability of a shift in any given direction, *p*_*i*_, is calculated as in Equation 1. The addition of +1 to the numerator ensures that *p*_*i*_>0, and the denominator is a normalisation so that ∑*p*_*i*_ = 1. As such, the equation entails that the probability of a move in any direction is determined by the difference between the fitness value after moving in that direction with step size *s*_*c*_ (i.e. *f*_*i*_) and the minimum fitness value possible across all mutations with step size of *s*_*c*_ (i.e. *f*_*min*_).

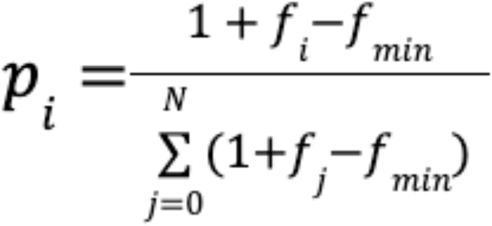

**Equation 1:**

Probability, for conservative mutations, of the mutational step being in any given direction.

## Results

Firstly, following up on the previously reported results whereby the standard genetic code appears to minimise differences in the average alternative-frame effect size for synonymous mutations, we confirmed that this property is very rare among alternative genetic codes. Among 10^7^ codes sharing the block structure of the standard genetic code, only 0.26% of codes performed equal or better to the standard genetic code in this property, measured as the standard deviation σ*D* between mutation effect values *D c* (Figure 3).

**Figure 3:**
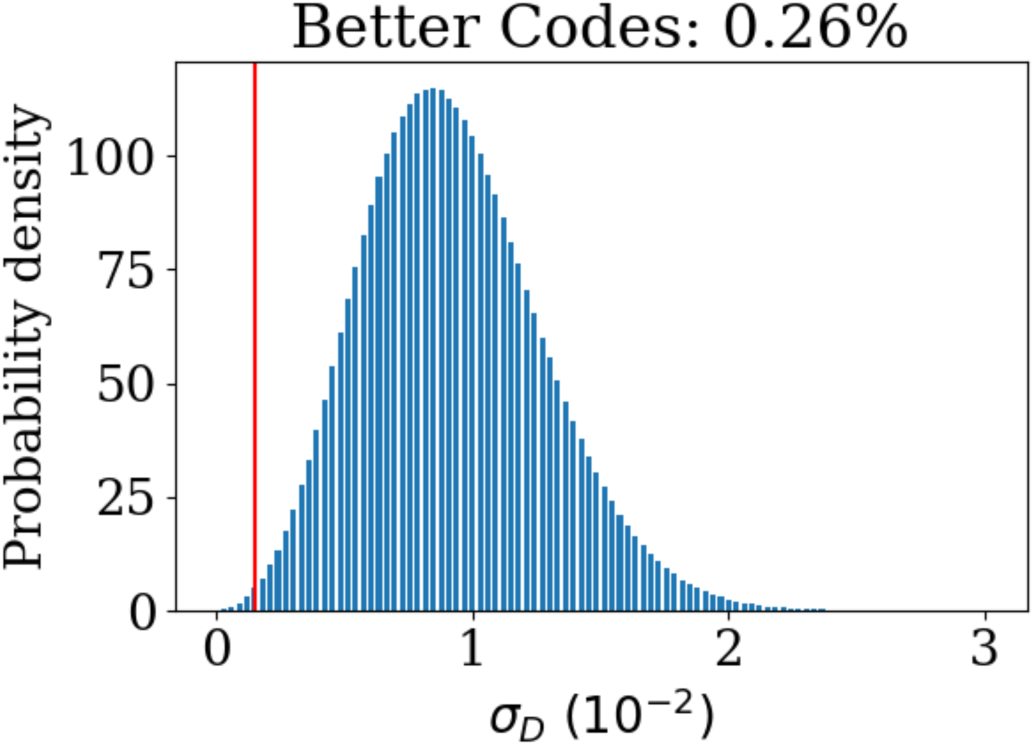
Distribution of standard deviations of mutation effect values across alternative frames (excluding frame “-1” i.e. sas13 in Figure 1). The position of the standard genetic code is shown with the vertical red line.

For the main part of this study we consider a simplification of the effects of large and small mutations in a rugged fitness landscape. Both evolvability and conservation are potentially advantageous for the evolution of overlapping genes. Evolvability allows exploring sequence space in order to find functional regions or improve fitness. Conservation allows functional sequences to be maintained in a population and not immediately degraded by new mutations, and conservative mutations also facilitate small-scale adaptation within a fitness peak. We hypothesise that for a given rugged fitness landscape, there is an optimal tradeoff between evolvability and robustness in mutation effects that will maximise the fitness of a population of novel sequences, over the long term. Whether the standard genetic code has actually achieved an optimal value is not addressed here - this study forms potential groundwork for further investigating this issue. In support of the hypothesis we present results from a simple model with two kinds of evolutionary processes, namely ‘evolvative’ and ‘conservative’, with sequences evolving on a fitness landscape with two broad fitness peaks of different heights, as described further in Methods.

The model illustrates the expected dynamics, whereby sequences accumulate in the larger fitness peak over time (Figure 4A). Eventually all sequences will be in the larger peak. Stochastic fluctuations in the proportion of sequences in the smaller fitness peak are larger than for the higher peak, reflecting the smaller peak’s lower capacity to retain sequences. If the probability of mutations being conservative, *pc*, is decreased then stochastic fluctuations in the distribution of sequences across peaks are larger, fewer sequences are found in the high peak at most time points, and the average fitness value of sequences is lower as many sequences are not in the high peak (Figure 4B). Average fitness over the medium to long term can thus be optimised by fine-tuning the degree of stochasticity (due to large-effect mutations) to a point where as many sequences as possible both reach and are retained in the higher peak.

**Figure 4:**
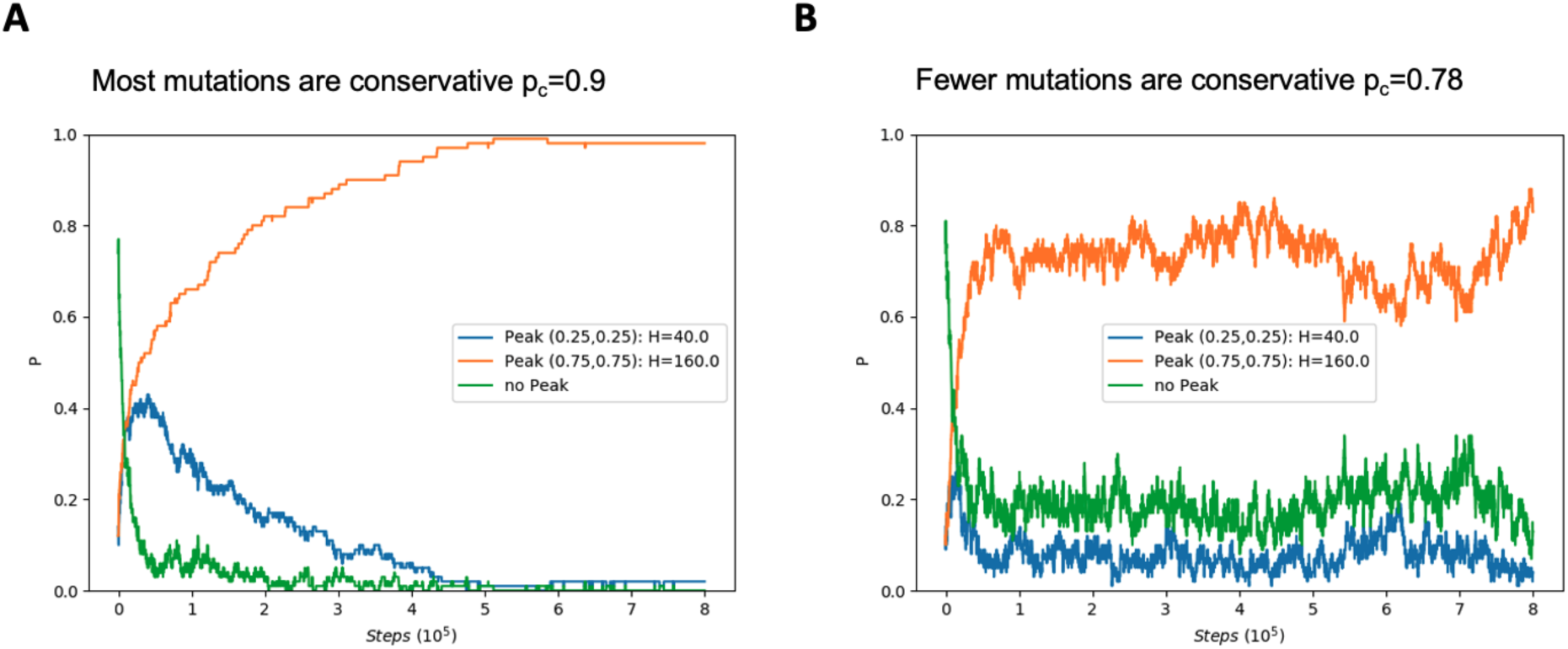
**A:** The proportion P of all sequences in one of the two peaks or outside any peak across 8·10^5^ mutation steps. Most sequences eventually end up in the higher peak (H=160) with some stochastic fluctuations. The data was created from 100 sequences, *sc* = 0.001, *se* = 0.01 and *pc* = 0.9. **B:** As for (A), but *pc* = 0.78. Sequences accumulate in the higher peak faster, but with larger stochastic fluctuations.

The model described so far uses two different step sizes, and calculates a ratio of how often each occurs. In order to use this model to test whether an average step size can be chosen so as to optimise sequence average fitness, the model is run to obtain average fitness values over a range of the parameters for step size and probability of each kind of mutation, as shown in Figure 5. Within the parameter space, we observe three qualitatively different regions, labelled as I, II, and III. In region I, conservative steps predominate and sequences remain in whichever fitness peak they are in, whether the high or low peak. In region III, the opposite tendency is observed and neither of the two peaks can retain sequences over the long term, i.e. stochastic fluctuations dominate the system. In the small high fitness area between I and III, i.e. region II, sequences are conserved in the higher but not the lower peak, as was observed in Figure 4A. The average mutation step size *<u>s</u>* is calculated with the equation *<u>s</u>* = *pc*.*sc* + (1 − *pc*)*se*. Fitting this equation (black line) to region II shows that *<u>s</u>* can be chosen so as to give a close approximation to the function which optimises fitness.

**Figure 5:**
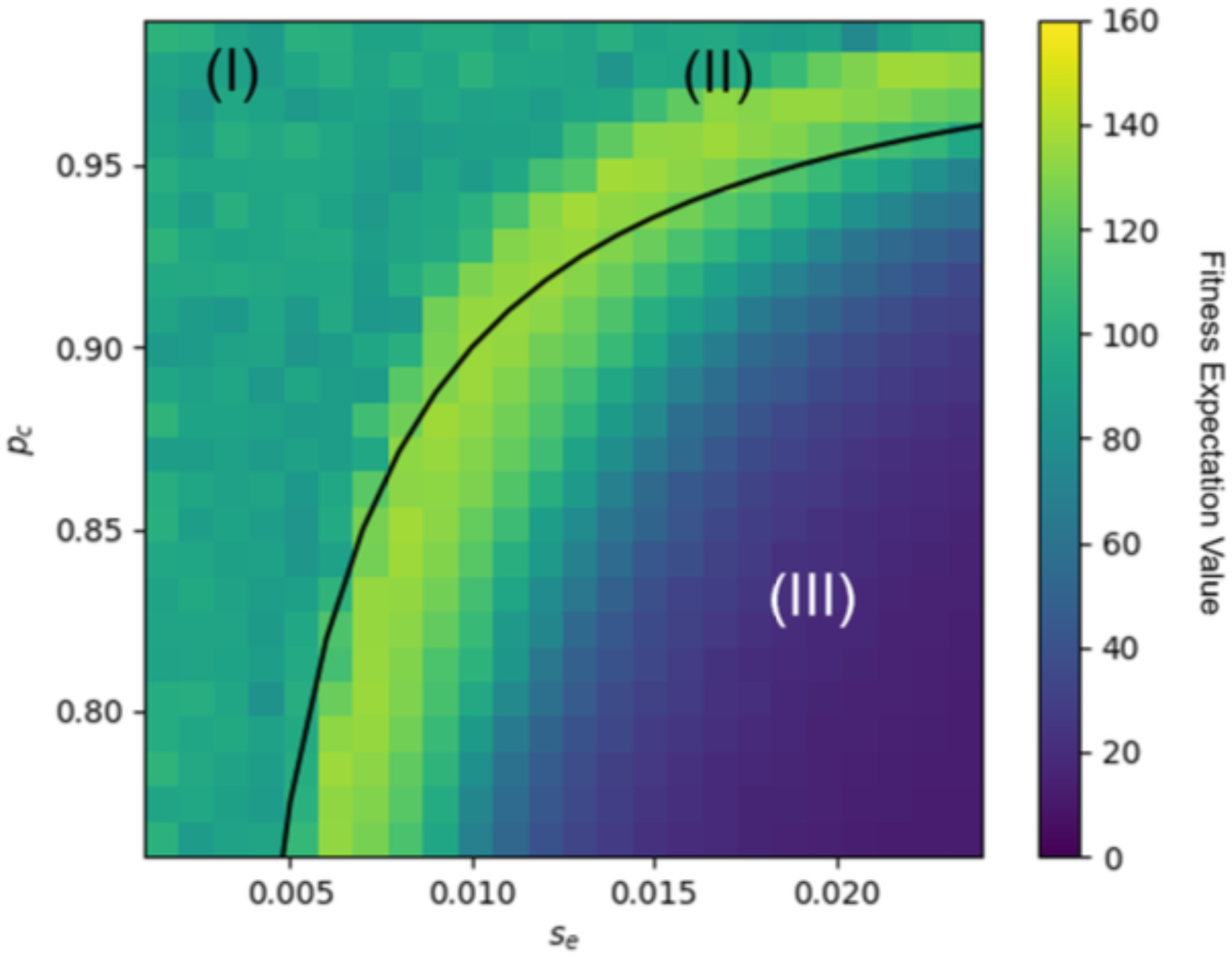
Average fitness value of 100 sequences after 8 · 10^5^ mutations for different values of conservative mutation probability *pc* and evolvative mutation step size *se*. The conservative mutation step size is fixed to *sc* = 0. 001. Fitness expectation values fall into three regions. In (I), sequences are retained in both peaks, in (II), sequences are restricted to the higher peak, and in (III), sequences are not retained in either peak. If we keep the average mutation effect size at a constant value (black line) we find almost the same functional relation between *pc* and *se* as that describing region (II) which optimises the expected fitness.

## Discussion

The core of this study is a model presented in order to examine the previously discovered property of consistent effect size across alternative reading frames resulting from synonymous changes in a reference frame. This model shows that the average mutation step size is able to be fine tuned so as to optimise the fitness of sequences on a given simple sequence landscape. It is only a toy model, presented as a proof of concept, laying groundwork for further investigation. Various aspects of the model can be called into question for lifelikeness. However, the key finding that the function for average step size so closely closely approximates the function for optimising average sequence fitness was surprising for us and appears to give real insight into evolutionary dynamics, albeit in a simplified form. After discussing ways in which this work can be developed or explored further, we will unpack some implications of the work and broader field of study of alternative reading frames.

Firstly, we note that the initial observation that led to model development adds another potential optimality to the growing list of interesting properties of the standard genetic code. Whether this is an artefact of some other property of the code or the method used for calculation, and how to interpret the finding if it does hold up both require further attention. It is important to note that some other putatively optimal properties have been shown to not be independently optimal and thus are perhaps an artefact of already-known facets of code structure. In addition to the putative frameshift optimality addressed above (Xu and Zhang 2021a), a prominent study in *Science* claimed that the code is near optimal for resource conservation (Shenhav and Zeevi 2020), but this claim has been subjected to rigorous critique in two response papers (Rozhoñová and Payne 2021; Xu and Zhang 2021b). The idea that the structure of the code may be a result of selection in general remains controversial (Massey 2008, 2016; Di Giulio 2018; Wichmann and Ardern 2019), but optimality is a distinct question from that of historical process.

Future research on this could include methodological improvements, more realistic modelling, investigation of OLG evolution in real biological data, and further investigation of the origins of the remarkable structure of the standard genetic code. Methodologically, it would be useful to clarify the relationship of the “minus one” (sas13) frame to the others - it appears likely that the approximately 20-fold difference in this relative frame is simply an artefact of the fact that this frame involves single codons matching to the reference frame rather than dicodons as for the other alternative frames, however this remains to be clarified. In any case, we believe that the results are striking enough whether or not this frame is included to be worthy of investigation. In terms of improving biological realism in the model, a number of steps can be taken. If the observed results hold across both more realistic models of evolving populations and diverse fitness landscapes this will support our claim - in principle we believe that they should. Finally regarding further developments, investigating sequence data from evolving OLGs is a potential new research field of its own. Until recently a large scale analysis was not possible, but improved methods for detecting overlapping genes’ protein expression such as ribosome profiling (Ingolia et al. 2009; Finkel et al. 2021) or detecting sequence features associated with OLGs (Firth 2014; Sealfon et al. 2015; Schlub, Buchmann, and Holmes 2018; Nelson, Ardern, and Wei 2020) will allow much larger sets for global studies across taxa.

The model can be seen as illustrating the process of finding function amidst the vast ‘hyperastronomical’ (Louis 2016) ocean of possible sequence space, the large majority of which is not functional even on relatively optimistic estimates (Keefe and Szostak 2001). Interpreted in this way, the model suggests that the right mutation effect size will help to maximise a population’s ability to find and retain new functions. Alternatively, instead of functionless regions of sequence space, the regions outside the peaks in the model can be conceived of as illustrating neutral networks across sequence space, with equivalent function. In either case, a particle moving into a peak represents finding functional novelty in sequence space. Whether evolvability or robustness is more important in real life depends on various parameters of the evolutionary processes in which alternative frame coding are involved. The idea of a trade-off between robustness and evolvability in the genetic code’s structure, facilitating the search for functional sequences, has been proposed before in the context of normal adaptive evolution (Tripathi and Deem 2018). Our results can be seen as an extension to this. It is also possible however that the apparent trade-off between robustness and evolvability in mutation effect size is better conceived of in another way than in terms of directly optimising fitness. For instance, as new sequences are sampled by evolution, small mutations allow a new sequence to be sampled relatively unchanged by stochastic translation or frameshift processes, while large mutations cause a ‘jump’ to new regions of sequence space; and both may be needed to optimally sample the total space and find new functions. As such, there could instead be a trade-off involving the time taken for sequence search, but we have not explored this here.

How does the potential origin of protein sequence novelty from alternative reading frames concretely impact biological reality? Recent findings on the role of alternative reading frames in protein origins is beginning to shed light on this. Alternative reading frames can potentially be used for protein novelty in at least three different ways: overprinting, remodelling, and frameshift mutations. The first to be discovered and investigated was the process of “overprinting”, where an out of frame open reading frame becomes translated and is retained due to some functional advantage. Conceivably the resulting overlapping gene pair could then be copied, and the homolog of the original gene then pseudogenized (Keese and Gibbs 1992). After some time this would leave essentially no trace of the original gene, and may explain the origin of some “orphan genes” - but to our knowledge no specific examples of this have yet been demonstrated. In eukaryotes, a related mechanism termed “mosaic translation” has been hypothesised but not demonstrated (Çakır et al. 2021) - this is where different parts of a spliced RNA transcript are read in different reading frames, producing “mosaic” proteins. The second process of remodelling has recently been demonstrated to play a non-trivial role in gene origins in *E. coli*. This is where gene fusion occurs between one or more frameshifted sequences (Watson, Lopez, and Bapteste 2022). A third process has also very recently been demonstrated, namely “mutually compensating frameshift mutations”, where two sequential frameshift mutations result in a partial frameshift which is tolerated in a protein sequence and opens up new sequence region for evolutionary exploration (Biba, Klink, and Bazykin 2022). More generally, if partially frameshifted sequences are tolerated they constitute new protein sequence space. All of these mechanisms bear some similarity to another process highlighted recently, where stochastic stop codon read-through allows some sequences after stop codons to contribute to protein novelty (Kosinski and Masel 2020).

Whether overprinting, remodelling, or frameshifting predominates is unclear, and the distribution is likely to be taxon dependent. In general, these processes have received little attention, likely at least partly due to a perceived limitation from evolutionary constraint (Yockey 1979; Lèbre and Gascuel 2017). Evolutionary constraint in OLGs has not yet been studied in much detail for specific gene pairs, apart from the case of overlapping genes in HIV-1. In this virus it has been shown firstly that constrained regions of overlapping genes in a pair are organised so as not to overlap (Fernandes et al. 2016) and secondly that even when domains overlap, functionally constrained residues in one protein are encoded overlapping codons which encode mutable codons in the other protein (Safari et al. 2021). Further, constraint can be conceived either negatively or positively. On the negative side, a study of constraints across reading frames concluded that the most constrained frame (the “minus two” frame, i.e. sas12 in Figure 1) is likely to be very rare in nature given the constraints in codons permitted in that frame (Lèbre and Gascuel 2017). On the other hand, constraint could bias overlapping frames towards functional sequences, acting as an approximate template. As described above, evidence for this actually being the case includes structure-promoting biases in both same-strand (Bartonek, Braun, and Zagrovic 2020) and antisense (Konecny et al. 1993) frames. It has already been shown that constraints from the structure of the genetic code facilitate Darwinian evolution (Firnberg and Ostermeier 2013), so it is reasonable to look for more cases of this.

In presenting the provocative hypothesis that this aspect of the structure of the standard genetic code may be “adaptive”, we are not making any specific claim about the processes behind the origin of the code’s structure. We have discussed some aspects of this in a previous publication (Wichmann and Ardern 2019). In this context “adaptive” (present usefulness) should not be conflated with “having been adapted” (historical process of fitting for a use), in the same way that “optimal for X” should not be conflated with “having been optimised for X”, and does not need to imply that it originated through a process of natural selection. Many aspects of biology are functional, may appear very “apt” for their functions, and can even be essential, without being a direct result primarily of selection, including RNA secondary structures (Dingle et al. 2022) and various examples of biochemical complexity (Schulz, Sendker, and Hochberg 2022). A related concept, although one which has accumulated a lot of conceptual baggage and related literature, is that of a spandrel (Gould and Lewontin 1979). The idea that the code has really been selected in order to be optimal across multiple parameters is perhaps implausible, given both the inherent difficulties in evolving any functional code (any change to the code will change multiple messages) and the limited timespan available for code evolution between the origin of life and the last universal common ancestor. The optimality of the code could be an example of what has been termed evolutionary inherency (Morris 2003), a widespread phenomenon where structures developed early in evolution end up being put to different functional use much later (such as seen in various components of animal nervous systems). Regardless of historical causes, the structure of the code and the uses it has in modern organisms are worth investigating. The main point we hope that the reader takes away from this study is the massive and exciting scope for further research regarding the contribution of the genetic code to evolvability and the origin of protein novelty from alternative reading frames.

## Acknowledgements

This manuscript is derived from work conducted as part of the PhD thesis of SW, recently submitted for examination at the Technical University of Munich and supervised by ZA. The model was presented in a poster at the conference “The Physics of Evolution” at the Francis Crick Institute, London, in July 2019. Thanks to everyone who gave feedback on the project, and to Siegfried Scherer and Klaus Neuhaus for supervision of the overall PhD and related projects.

## Notes

### Competing Interest Statement

The authors have declared no competing interest.

## References

Affram, Yvonne, Juan C. Zapata, Zahra Gholizadeh, William D. Tolbert, Wei Zhou, Maria D. Iglesias-Ussel, Marzena Pazgier, Krishanu Ray, Olga S. Latinovic, and Fabio Romerio. 2019. “The HIV-1 Antisense Protein ASP Is a Transmembrane Protein of the Cell Surface and an Integral Protein of the Viral Envelope.” Journal of Virology 93 (21). https://doi.org/10.1128/JVI.00574-19.

Barrell, B. G., G. M. Air, and C. A. Hutchison. 1976. “Overlapping Genes in Bacteriophage ΦX174.” Nature 264 (5581): 34–41.

Bartonek, Lukas, Daniel Braun, and Bojan Zagrovic. 2020. “Frameshifting Preserves Key Physicochemical Properties of Proteins.” Proceedings of the National Academy of Sciences of the United States of America 117 (11): 5907–12.

Biba, Dmitry, Galya Klink, and Georgii Bazykin. 2022. “Pairs of Mutually Compensatory Frameshifting Mutations Contribute to Protein Evolution.” Molecular Biology and Evolution, February. https://doi.org/10.1093/molbev/msac031.

Brandes, Nadav, and Michal Linial. 2016. “Gene Overlapping and Size Constraints in the Viral World.” Biology Direct 11 (May): 26.

Buhrman, Harry, Peter T. S. van der Gulik, Steven M. Kelk, Wouter M. Koolen, and Leen Stougie. 2011. “Some Mathematical Refinements Concerning Error Minimization in the Genetic Code.” IEEE/ACM Transactions on Computational Biology and Bioinformatics / IEEE, ACM 8 (5): 1358–72.

Çakır, Umut, Noujoud Gabed, Marie Brunet, Xavier Roucou, and Igor Kryvoruchko. 2021. “Mosaic Translation Hypothesis: Chimeric Polypeptides Produced via Multiple Ribosomal Frameshifting as a Basis for Adaptability.” The FEBS Journal, November. https://doi.org/10.1111/febs.16269.

Cao, Xiongwen, Alexandra Khitun, Yang Luo, Zhenkun Na, Thitima Phoodokmai, Khomkrit Sappakhaw, Elizabeth Olatunji, Chayasith Uttamapinant, and Sarah A. Slavoff. 2021. “Alt-RPL36 Downregulates the PI3K-AKT-mTOR Signaling Pathway by Interacting with TMEM24.” Nature Communications 12 (1): 508.

Carter, Charles W., Jr. 2021. “Simultaneous Codon Usage, the Origin of the Proteome, and the Emergence of de-Novo Proteins.” Current Opinion in Structural Biology 68 (June): 142–48.

Cassan, Elodie, Anne-Muriel Arigon-Chifolleau, Jean-Michel Mesnard, Antoine Gross, and Olivier Gascuel. 2016. “Concomitant Emergence of the Antisense Protein Gene of HIV-1 and of the Pandemic.” Proceedings of the National Academy of Sciences of the United States of America 113 (41): 11537–42.

Chen, J. Z., D. M. Fowler, and N. Tokuriki. 2022. “Environmental Selection and Epistasis in an Empirical Phenotype-Environment-Fitness Landscape.” Nature Ecology & Evolution, February. https://doi.org/10.1038/s41559-022-01675-5.

Di Giulio, Massimo. 2018. “A Non-Neutral Origin for Error Minimization in the Origin of the Genetic Code.” Journal of Molecular Evolution 86 (9): 593–97.

Dingle, Kamaludin, Fatme Ghaddar, Petr Šulc, and Ard A. Louis. 2022. “Phenotype Bias Determines How Natural RNA Structures Occupy the Morphospace of All Possible Shapes.” Molecular Biology and Evolution 39 (1). https://doi.org/10.1093/molbev/msab280.

Fernandes, Jason D., Tyler B. Faust, Nicolas B. Strauli, Cynthia Smith, David C. Crosby, Robert L. Nakamura, Ryan D. Hernandez, and Alan D. Frankel. 2016. “Functional Segregation of Overlapping Genes in HIV.” Cell 167 (7): 1762–73.e12.

Finkel, Yaara, Orel Mizrahi, Aharon Nachshon, Shira Weingarten-Gabbay, David Morgenstern, Yfat Yahalom-Ronen, Hadas Tamir, et al. 2021. “The Coding Capacity of SARS-CoV-2.” Nature 589 (7840): 125–30.

Firnberg, Elad, and Marc Ostermeier. 2013. “The Genetic Code Constrains yet Facilitates Darwinian Evolution.” Nucleic Acids Research 41 (15): 7420–28.

Firth, Andrew E. 2014. “Mapping Overlapping Functional Elements Embedded within the Protein-Coding Regions of RNA Viruses.” Nucleic Acids Research 42 (20): 12425–39.

Firth, Andrew E. 2020. “A Putative New SARS-CoV Protein, 3c, Encoded in an ORF Overlapping ORF3a.” The Journal of General Virology 101 (10): 1085–89.

Firth, Andrew E., and Ian Brierley. 2012. “Non-Canonical Translation in RNA Viruses.” The Journal of General Virology 93 (Pt 7): 1385–1409.

Fisher, Ronald A. 1930. “The Genetical Theory of Natural Selection. Clarendon.” Oxford. https://doi.org/10.5962/bhl.title.27468.

Freeland, S. J., and L. D. Hurst. 1998. “The Genetic Code Is One in a Million.” Journal of Molecular Evolution 47 (3): 238–48.

Freeland, S. J., R. D. Knight, L. F. Landweber, and L. D. Hurst. 2000. “Early Fixation of an Optimal Genetic Code.” Molecular Biology and Evolution 17 (4): 511–18.

Freeland, Stephen J. 2002. Genetic Programming and Evolvable Machines 3 (2): 113–27.

Gelsinger, Diego Rivera, Emma Dallon, Rahul Reddy, Fuad Mohammad, Allen R. Buskirk, and Jocelyne DiRuggiero. 2020. “Ribosome Profiling in Archaea Reveals Leaderless Translation, Novel Translational Initiation Sites, and Ribosome Pausing at Single Codon Resolution.” Nucleic Acids Research 48 (10): 5201–16.

Gould, Stephen Jay, and Richard C. Lewontin. 1979. “5 The Spandrels of San Marco and the Panglossian Paradigm: A Critique of the Adaptationist Programme.” Conceptual Issues in Evolutionary Biology 205: 79.

Hücker, Sarah M., Sonja Vanderhaeghen, Isabel Abellan-Schneyder, Siegfried Scherer, and Klaus Neuhaus. 2018. “The Novel Anaerobiosis-Responsive Overlapping Gene Ano Is Overlapping Antisense to the Annotated Gene ECs2385 of Escherichia Coli O157:H7 Sakai.” Frontiers in Microbiology 9 (May): 931.

Hücker, Sarah M., Sonja Vanderhaeghen, Isabel Abellan-Schneyder, Romy Wecko, Svenja Simon, Siegfried Scherer, and Klaus Neuhaus. 2018. “A Novel Short L-Arginine Responsive Protein-Coding Gene (laoB) Antiparallel Overlapping to a CadC-like Transcriptional Regulator in Escherichia Coli O157:H7 Sakai Originated by Overprinting.” BMC Evolutionary Biology 18 (1): 21.

Ilardo, Melissa, Rudrarup Bose, Markus Meringer, Bakhtiyor Rasulev, Natalie Grefenstette, James Stephenson, Stephen Freeland, Richard J. Gillams, Christopher J. Butch, and H. James Cleaves 2nd. 2019. “Adaptive Properties of the Genetically Encoded Amino Acid Alphabet Are Inherited from Its Subsets.” Scientific Reports 9 (1): 12468.

Ilardo, Melissa, Markus Meringer, Stephen Freeland, Bakhtiyor Rasulev, and H. James Cleaves 2nd. 2015. “Extraordinarily Adaptive Properties of the Genetically Encoded Amino Acids.” Scientific Reports 5 (March): 9414.

Ingolia, Nicholas T., Sina Ghaemmaghami, John R. S. Newman, and Jonathan S. Weissman. 2009. “Genome-Wide Analysis in Vivo of Translation with Nucleotide Resolution Using Ribosome Profiling.” Science 324 (5924): 218–23.

Itzkovitz, Shalev, and Uri Alon. 2007. “The Genetic Code Is Nearly Optimal for Allowing Additional Information within Protein-Coding Sequences.” Genome Research 17 (4): 405–12.

Keefe, A. D., and J. W. Szostak. 2001. “Functional Proteins from a Random-Sequence Library.” Nature 410 (6829): 715–18.

Keese, P. K., and A. Gibbs. 1992. “Origins of Genes:’ Big Bang’ or Continuous Creation?” Of the National Academy of Sciences. https://www.pnas.org/content/89/20/9489.short.

Khan, Yousuf A., Irwin Jungreis, James C. Wright, Jonathan M. Mudge, Jyoti S. Choudhary, Andrew E. Firth, and Manolis Kellis. 2020. “Evidence for a Novel Overlapping Coding Sequence in POLG Initiated at a CUG Start Codon.” BMC Genetics 21 (1): 25.

Kolata, G. B. 1977. “Overlapping Genes: More than Anomalies?” Science 196 (4295): 1187–88.

Konecny, J., M. Eckert, M. Schöniger, and G. L. Hofacker. 1993. “Neutral Adaptation of the Genetic Code to Double-Strand Coding.” Journal of Molecular Evolution 36 (5): 407–16.

Kosinski, Luke J., and Joanna Masel. 2020. “Readthrough Errors Purge Deleterious Cryptic Sequences, Facilitating the Birth of Coding Sequences.” Molecular Biology and Evolution 37 (6): 1761–74.

Kreitmeier, Michaela, Zachary Ardern, Miriam Abele, Christina Ludwig, Siegfried Scherer, and Klaus Neuhaus. 2022. “Spotlight on Alternative Frame Coding: Two Long Overlapping Genes in Pseudomonas Aeruginosa Are Translated and under Purifying Selection.” iScience 25 (2): 103844.

Lèbre, Sophie, and Olivier Gascuel. 2017. “The Combinatorics of Overlapping Genes.” Journal of Theoretical Biology 415 (February): 90–101.

Loughran, Gary, Alexander V. Zhdanov, Maria S. Mikhaylova, Fedor N. Rozov, Petr N. Datskevich, Sergey I. Kovalchuk, Marina V. Serebryakova, et al. 2020. “Unusually Efficient CUG Initiation of an Overlapping Reading Frame in POLG mRNA Yields Novel Protein POLGARF.” Proceedings of the National Academy of Sciences of the United States of America 117 (40): 24936–46.

Louis, Ard A. 2016. “Contingency, Convergence and Hyper-Astronomical Numbers in Biological Evolution.” Studies in History and Philosophy of Biological and Biomedical Sciences 58 (August): 107–16.

Massey, Steven E. 2008. “A Neutral Origin for Error Minimization in the Genetic Code.” Journal of Molecular Evolution 67 (5): 510–16.

Massey, Steven E. 2016. “The Neutral Emergence of Error Minimized Genetic Codes Superior to the Standard Genetic Code.” Journal of Theoretical Biology 408 (November): 237–42.

Mayer-Bacon, Christopher, and Stephen J. Freeland. 2021. “A Broader Context for Understanding Amino Acid Alphabet Optimality.” Journal of Theoretical Biology 520 (July): 110661.

Meydan, Sezen, Nora Vázquez-Laslop, and Alexander S. Mankin. 2018. “Genes within Genes in Bacterial Genomes.” Microbiology Spectrum 6 (4). https://doi.org/10.1128/microbiolspec.RWR-0020-2018.

Miyata, T., and T. Yasunaga. 1978. “Evolution of Overlapping Genes.” Nature 272 (5653): 532–35.

Morris, Simon Conway. 2003. “Life’s Solution.” https://doi.org/10.1017/cbo9780511535499.

Mudge, Jonathan M., Jorge Ruiz-Orera, John R. Prensner, Marie A. Brunet, Jose Manuel Gonzalez, Michele Magrane, Thomas Martinez, et al. 2021. “A Community-Driven Roadmap to Advance Research on Translated Open Reading Frames Detected by Ribo-Seq.” bioRxiv. https://doi.org/10.1101/2021.06.10.447896.

Nelson, Chase W., Zachary Ardern, Tony L. Goldberg, Chen Meng, Chen-Hao Kuo, Christina Ludwig, Sergios-Orestis Kolokotronis, and Xinzhu Wei. 2020. “Dynamically Evolving Novel Overlapping Gene as a Factor in the SARS-CoV-2 Pandemic.” eLife 9 (October). https://doi.org/10.7554/eLife.59633.

Nelson, Chase W., Zachary Ardern, and Xinzhu Wei. 2020. “OLGenie: Estimating Natural Selection to Predict Functional Overlapping Genes.” Molecular Biology and Evolution 37 (8): 2440–49.

Ohno, Susumu. 1970. Evolution by Gene Duplication. Springer Berlin Heidelberg.

Osawa, S. 1995. Evolution of the Genetic Code. Oxford University Press.

Payne, Joshua L., and Andreas Wagner. 2019. “The Causes of Evolvability and Their Evolution.” Nature Reviews. Genetics 20 (1): 24–38.

Richter, Hendrik, and Andries Engelbrecht, eds. 2014. Recent Advances in the Theory and Application of Fitness Landscapes. Springer, Berlin, Heidelberg.

Rozhoňová, Hana, and Joshua L. Payne. 2021. “Little Evidence the Standard Genetic Code Is Optimized for Resource Conservation.” Molecular Biology and Evolution 38 (11): 5127–33.

Safari, Maliheh, Bhargavi Jayaraman, Shumin Yang, Cynthia Smith, Jason D. Fernandes, and Alan D. Frankel. 2021. “Functional and Structural Segregation of Overlapping Helices in HIV-1.” bioRxiv. https://doi.org/10.1101/2021.07.15.452440.

Schlub, Timothy E., Jan P. Buchmann, and Edward C. Holmes. 2018. “A Simple Method to Detect Candidate Overlapping Genes in Viruses Using Single Genome Sequences.” Molecular Biology and Evolution 35 (10): 2572–81.

Schulz, Luca, Franziska L. Sendker, and Georg K. A. Hochberg. 2022. “Non-Adaptive Complexity and Biochemical Function.” Current Opinion in Structural Biology 73 (April): 102339.

Sealfon, Rachel S., Michael F. Lin, Irwin Jungreis, Maxim Y. Wolf, Manolis Kellis, and Pardis C. Sabeti. 2015. “FRESCo: Finding Regions of Excess Synonymous Constraint in Diverse Viruses.” Genome Biology 16 (February): 38.

Shenhav, Liat, and David Zeevi. 2020. “Resource Conservation Manifests in the Genetic Code.” Science 370 (6517): 683–87.

Siegel, Andrew F., and Walter M. Fitch. 1980. “Degeneracy When DNA Codes for Overlapping Genes.” Mathematical Biosciences 49 (1): 1–16.

Smith, T. F., and M. S. Waterman. 1981. “Overlapping Genes and Information Theory.” Journal of Theoretical Biology 91 (2): 379–80.

Szekely, M. 1978. “Triple Overlapping Genes.” Nature 272 (5653): 492.

Tenaillon, O. 2014. “The Utility of Fisher’s Geometric Model in Evolutionary Genetics.” Annual Review of Ecology, Evolution, and Systematics 45 (November): 179–201.

Tripathi, Shubham, and Michael W. Deem. 2018. “The Standard Genetic Code Facilitates Exploration of the Space of Functional Nucleotide Sequences.” Journal of Molecular Evolution 86 (6): 325–39.

Vakirlis, Nikolaos, Anne-Ruxandra Carvunis, and Aoife McLysaght. 2020. “Synteny-Based Analyses Indicate That Sequence Divergence Is Not the Main Source of Orphan Genes.” eLife 9 (February). https://doi.org/10.7554/eLife.53500.

Vanderhaeghen, Sonja, Barbara Zehentner, Siegfried Scherer, Klaus Neuhaus, and Zachary Ardern. 2018. “The Novel EHEC Gene Asa Overlaps the TEGT Transporter Gene in Antisense and Is Regulated by NaCl and Growth Phase.” Scientific Reports 8 (1): 17875.

Visser, J. Arjan G. M. de, and Joachim Krug. 2014. “Empirical Fitness Landscapes and the Predictability of Evolution.” Nature Reviews. Genetics 15 (7): 480–90.

Watson, Andrew K., Philippe Lopez, and Eric Bapteste. 2022. “Hundreds of Out-of-Frame Remodeled Gene Families in the Escherichia Coli Pangenome.” Molecular Biology and Evolution 39 (1). https://doi.org/10.1093/molbev/msab329.

Wei, Xinzhu, and Jianzhi Zhang. 2015. “A Simple Method for Estimating the Strength of Natural Selection on Overlapping Genes.” Genome Biology and Evolution. https://doi.org/10.1093/gbe/evu294.

Wichmann, Stefan, and Zachary Ardern. 2019. “Optimality in the Standard Genetic Code Is Robust with Respect to Comparison Code Sets.” Bio Systems 185 (November): 104023.

Willis, Sara, and Joanna Masel. 2018. “Gene Birth Contributes to Structural Disorder Encoded by Overlapping Genes.” Genetics 210 (1): 303–13.

Wright, Bradley W., Mark P. Molloy, and Paul R. Jaschke. 2022. “Overlapping Genes in Natural and Engineered Genomes.” Nature Reviews. Genetics 23 (3): 154–68.

Wright, Bradley W., Zixin Yi, Jonathan S. Weissman, and Jin Chen. 2022. “The Dark Proteome: Translation from Noncanonical Open Reading Frames.” Trends in Cell Biology 32 (3): 243–58.

Xu, Haiqing, and Jianzhi Zhang. 2021a. “On the Origin of Frameshift-Robustness of the Standard Genetic Code.” Molecular Biology and Evolution 38 (10): 4301–9.

Xu, Haiqing, and Jianzhi Zhang. 2021b. “Is the Genetic Code Optimized for Resource Conservation?” Molecular Biology and Evolution 38 (11): 5122–26.

Yockey, H. P. 1979. “Do Overlapping Genes Violate Molecular Biology and the Theory of Evolution?” Journal of Theoretical Biology 80 (1): 21–26.

Yockey, H. P. 1981. “Rebuttal of ‘overlapping Genes and Information Theory.’” Journal of Theoretical Biology 91 (2): 381–82.

Zehentner, Barbara, Zachary Ardern, Michaela Kreitmeier, Siegfried Scherer, and Klaus Neuhaus. 2020a. “A Novel pH-Regulated, Unusual 603 Bp Overlapping Protein Coding Gene Pop Is Encoded Antisense to ompA in Escherichia Coli O157:H7 (EHEC).” Frontiers in Microbiology 11 (March): 377.

Zehentner, Barbara, Zachary Ardern, Michaela Kreitmeier, Siegfried Scherer, and Klaus Neuhaus. 2020b. “Evidence for Numerous Embedded Antisense Overlapping Genes in Diverse E. Coli Strains.” bioRxiv. https://doi.org/10.1101/2020.11.18.388249.

Zhu, Wen, and Stephen Freeland. 2006. “The Standard Genetic Code Enhances Adaptive Evolution of Proteins.” Journal of Theoretical Biology 239 (1): 63–70.

Zull, J. E., and S. K. Smith. 1990. “Is Genetic Code Redundancy Related to Retention of Structural Information in Both DNA Strands?” Trends in Biochemical Sciences 15 (7): 257–61.

